# Heterozygosity for Crohn’s Disease Risk Allele of *ATG16L1* Protects against Bacterial Infection

**DOI:** 10.1101/2024.02.22.581423

**Authors:** Xiaomin Yao, Eugene Rudensky, Patricia K. Martin, Erin E. Zwack, Zhengxiang He, Glaucia C. Furtado, Sérgio A. Lira, Victor J. Torres, Bo Shopsin, Ken Cadwell

## Abstract

The T300A substitution in ATG16L1 associated with Crohn’s Disease impairs autophagy, yet up to 50% of humans are heterozygous for this allele. Here we demonstrate that heterozygosity for the analogous substitution in mice (*Atg16L1^T316A^*), but not homozygosity, protects against lethal *Salmonella enterica* Typhimurium infection. One copy of *Atg16L1^T316A^* was sufficient to enhance cytokine production through inflammasome activation, which was necessary for protection. In contrast, two copies of *Atg16L1^T316A^* inhibited the autophagy-related process of LC3-associated phagocytosis (LAP) and increased susceptibility. Macrophages from human donors heterozygous for *ATG16L1^T300A^*displayed elevated inflammasome activation while homozygosity impaired LAP, similar to mice. These results clarify how the T300A substitution impacts ATG16L1 function and suggest it can be beneficial to heterozygous carriers, providing an explanation for its prevalence.

**One-Sentence Summary:** Heterozygosity of Crohn’s diseases risk variant *ATG16L1 T300A* confers protection against bacterial infections.

## Main Text

Crohn’s Disease (CD) is an inflammatory bowel disease characterized by recurring and relapsing inflammation of the gastrointestinal tract with a complex etiology involving both genetic susceptibility and environmental factors, such as enteric infection and microbiota alterations (*1, 2*). A variant of the autophagy gene *ATG16L1* (rs2241880) that causes a threonine to alanine substitution at position 300 (T300A) is among the highest genetic risk factors for developing CD, however, over 50% of the global population, and over 70% of certain sub-populations, carry at least one copy of the allele (*3*). During autophagy, ATG16L1 mediates lipidation of ATG8 family members (LC3 and GABARAP proteins) to generate double-membraned autophagosomes that can fuse with the lysosome, leading to degradation and recycling of cytosolic cargoes (*4*). ATG8 lipidation by ATG16L1 is also required for autophagosome-independent processes, such as membrane repair and cargo trafficking during phagocytosis and exocytosis, collectively referred to as Conjugation of Atg8 to Single Membranes (CASM) (*5, 6*). The T300A substitution (T316A in mice) can impair autophagy and CASM through either destabilizing ATG16L1 protein or disrupting interactions with binding partners (*7–9*). Consistent with the link to an immune-mediated intestinal disorder, mice homozygous for knock-in alleles encoding ATG16L1^T316A^ display abnormalities in the intestinal epithelium, microbiota composition, and cytokine responses to infections (*8–10*).

Autophagy features prominently during *Salmonella* infection. *Salmonella* species represent food-borne pathogens, causing millions of illnesses ranging from self-limiting gastroenteritis to sepsis and death. *Salmonella enterica* serovar Typhimurium (herein *Salmonella*) has a broad host range and has been used to model typhoid-like disease in mice (*11*). Autophagy can restrict internalized *Salmonella* in cultured mammalian cells or support bacterial replication by maintaining the membrane integrity of the intracellular replicative niche known as the *Salmonella* containing vacuole (SCV) (*12–16*). Deletion of autophagy genes in the intestinal epithelium or myeloid compartment renders mice more susceptible to *Salmonella* infection, and a small molecule agonist of autophagy promotes defense *in vivo* (*17–19*). Additionally, *Salmonella* produces several effector molecules that alter the autophagy pathway, including SopF that inhibits ATG16L1 (*20*). Thus, *Salmonella* has adapted to their mammalian host by acquiring strategies to evade and subvert the autophagy pathway.

This role of autophagy revealed using *Salmonella* as a model pathogen suggests that loss-of-function due to homozygosity for the *ATG16L1^T300A^*variant contributes to disease through disruption of innate immunity. However, the plurality of the humans carries only one copy of the disease associated allele. Thus, we examined the impact of *ATG16L1^T300A^* on *Salmonella* infection with a focus on heterozygosity.

## Results

### *Atg16L1^T316A^* heterozygosity confers resistance to bacterial infection

To assess the impact of the CD risk variant on antibacterial defense *in vivo*, we orally inoculated littermate *Atg16L1* wild-type (*Atg16L1*^T/T^), *Atg16L1^T316A^* heterozygous (*Atg16L1*^T/A^), and *Atg16L1^T316A^* homozygous (*Atg16L1*^A/A^) knock-in mice with *Salmonella*. The time course of lethality was modestly but significantly accelerated in *Atg16L1*^A/A^ mice compared with *Atg16L1*^T/T^ mice (Fig. 1A), confirming previous findings demonstrating that T300A homozygosity increases susceptibility to bacterial pathogens (*8, 9, 21*). In stark contrast to the complete lethality observed in these mice homozygous for either the wild-type or T316A alleles, nearly all *Atg16L1*^T/A^ animals survived the infection (Fig. 1A). The three genotypes of mice had similar levels of *Salmonella* burden in stool at the early time points. On day 7 when the genotypes began to display differences in survival, the numbers of bacteria shed by *Atg16L1^T/A^*mice was reduced compared with *Atg16L1^T/T^* and *Atg16L1^A/A^*mice (Fig. 1B). Similarly, *Atg16L1^T/A^* mice displayed substantially lower bacterial counts in the filtering organs (Fig. 1C).

**Fig. 1.**
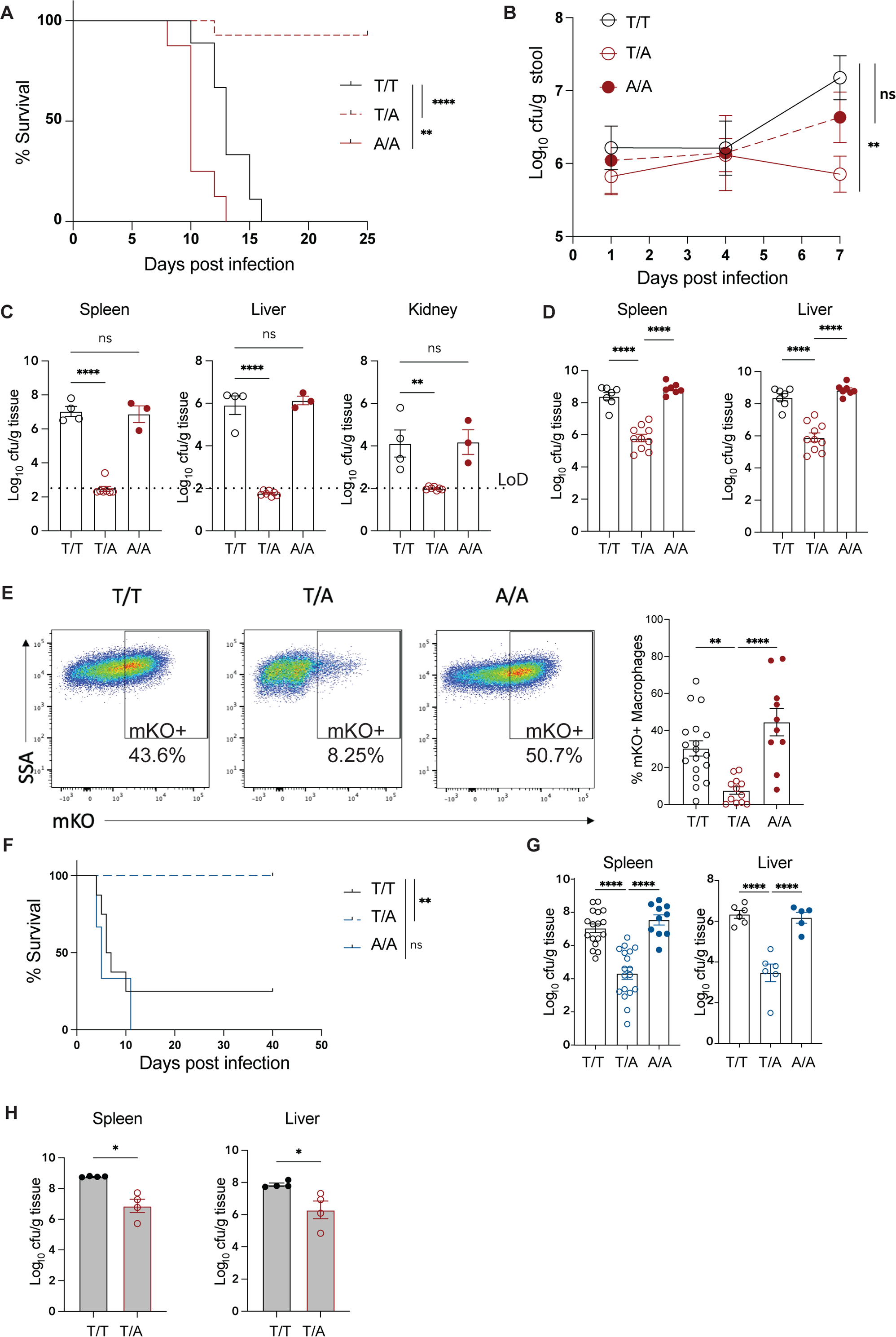
*Atg16L1 T316A* heterozygous mice display enhanced resistance to bacterial infection. (**A**) Wild-type (T/T, n=9) mice and mice heterozygous (T/A, n=14) or homozygous (A/A, n=8) for the *Atg16L1 T316A* variant were inoculated with 2×10^4^ colony forming units (cfu) of *Salmonella* by oral gavage and monitored for survival. (**B**) *Salmonella* cfu from stool of mice on indicated days post-inoculation as in (A). T/T, n=10; T/A, n=11; A/A, n=15. (**C**) *Salmonella* cfu in the liver, kidney, and spleen on day 7 post-inoculation, LoD, limit of detection. T/T, n=4; T/A, n=7; A/A, n=3. (**D**) Bacterial burden in indicated organs 2 days post intraperitoneal (IP) injection with 2×10^5^ cfu *Salmonella*. T/T, n=7; T/A, n=10; A/A, n=7. (**E**) Mice were injected IP as in (D) with mKO-expressing *Salmonella* and peritoneal exudate cells were analyzed by FACS for intracellular bacteria after 2 days. Gating on live, CD45^+^, F4/80^+^, CD11b^+^ cells for macrophages. T/T, n=18; T/A, n=12; A/A, n=10. (**F**) Survival of mice following IP inoculation with 2×10^4^ cfu *L. monocytogenes*. T/T, n=7; T/A, n=8; A/A, n=3. (**G**) *L. monocytogenes* cfu in spleen and liver of mice 3 days post IP inoculation as in (F). For spleen T/T, n=17; T/A, n=18; A/A, n=10; for liver T/T, n=6; T/A, n=6; A/A, n=5. (**H**) Chimeras generated by reconstituting CD45.1 mice with bone marrow of indicated genotypes from an independently generated strain of *Atg16L1^T316A^* mice were infected as in D and bacterial burden in filtering organs was determined 2 days post infection, n=4 each. Data points in (B, C, D, E, G, H and I) represent individual mice and bars represent mean ± SEM. All data were collected from at least two independent experiments. ^∗^p < 0.05; ^∗∗^p < 0.01; ^∗∗∗^p < 0.001; ^∗∗∗∗^p < 0.0001; ns, not significant, by log-rank Mantel-Cox test in (A) and (F), one-way ANOVA with Holm-Sidak test for (B, C, D, E, G) and Mann-Whitney test for (H).

Bacterial burden in *Atg16L1*^T/A^ mice was reduced when we inoculated mice by intraperitoneal (IP) injection (Fig. 1D), which bypasses defense mechanisms at the intestinal barrier that precede dissemination. During systemic infection, *Salmonella* preferentially replicates in myeloid cells, accumulating in tissues enriched for macrophages (*22–25*). We inoculated mice using *Salmonella* expressing the fluorophore mKO to identify infected cells by flow cytometry. Two days post IP inoculation, when the peritoneal cavity is enriched for myeloid cells (Fig. S1A), we found that the proportion of mKO+ cells was reduced in *Atg16L1*^T/A^ mice compared with *Atg16L1*^A/A^ and *Atg16L1*^T/T^ controls, especially in macrophages (Fig. 1E and S1B). To test whether the protection conferred by *Atg16L1^T316A^*heterozygosity was specific to *Salmonella*, we infected mice with *Listeria monocytogenes*, another bacterial pathogen that infects macrophages and uses multiple strategies to evade or subvert autophagy proteins to establish successful infection (*26–32*). Again, *Atg16L1*^T/A^ survived IP inoculation with a dose of *L. monocytogenes* that was lethal to *Atg16L1*^T/T^ and *Atg16L1*^A/A^ mice, with a corresponding reduction in bacterial burden in organs (Fig. 1F-G). Therefore, ATG16L1 T316A heterozygosity offers an unexpected advantage to the host by providing profound protection from two well-characterized model pathogens.

To facilitate downstream experiments, we generated chimeras in which irradiated CD45.1+ mice received bone marrow (BM) cells from *Atg16LT^T/T^* and *Atg16L1^T/A^* mice, which led to >95% of the cells in the peritoneal cavity to be of CD45.2+ donor origin (Fig. S1C). Mice reconstituted with *Atg16L1^T/A^* BM displayed a reduction in *Salmonella* burden following IP injection compared with those receiving *Atg16L1^T/T^* BM (Fig. S1D), indicating that *Atg16L1 T316A* heterozygosity in the hematopoietic compartment is sufficient for resistance. Given the unusually strong and specific effect of heterozygosity, we validated our findings using an independently generated *Atg16L1^T316A^*knock-in mouse line (see methods). Our ability to utilize BM chimeras allowed us to examine *Atg16L1^T316A^* mice generated at another institution without the need to transfer live animals. Similar to the above experiment, chimeric animals reconstituted with *Atg16L1^T/A^* BM from these independently generated mice displayed less bacteria in organs compared with those reconstituted with matched *Atg16L1^T/T^*BM (Fig. 1I and Fig. S1E).

### *Atg16L1^T316A^* heterozygosity is sufficient to enhance proinflammatory responses at the transcriptional level

To determine whether the reduction in the proportion of macrophages infected by *Salmonella* in *Atg16L1^T/A^* mice (Fig. 1E) could be due to superior control of internalized bacteria, we infected BM-derived macrophages (BMDMs) from *Atg16L1^T/T^*, *Atg16L1^T/A^,* or *Atg16L1^A/A^* mice *in vitro*. BMDMs of all three genotypes demonstrated equal uptake of *Salmonella* and bacterial burden remained similar between groups at later time points (Fig. S2A). Thus, the *in vivo* protection conferred by *Atg16L1^T316A^* heterozygosity might not be due to cell autonomous control of infection. To further examine this possibility, we carried out a mixed BM chimera experiment in which BM cells from *Atg16L1^T316A^*mice (on CD45.2 background) were mixed with BM cells from congenic CD45.1 wildtype mice at a 1:1 ratio, and then used to reconstitute GFP+ wildtype recipients. This experimental set up allows comparison of donor-derived *Atg16L1* mutant and wildtype cells infected with *Salmonella* (mKO+) within the same host, which can be distinguished from residual recipient macrophages expressing GFP. As expected, similar proportions of *Atg16L1^T/T^*CD45.2+donor cells were infected by *Salmonella* as the wildtype CD45.1+ cells. *Atg16L1^A/A^* and *Atg16L1^T/A^* CD45.2+ donor cells were also similarly infected as wildtype CD45.1+ cells, indicating that the protective effect of *Atg16L1^T316A^*heterozygosity was hindered by the presence of wildtype cells (Fig. S2B). Additionally, we found the proportion of *Salmonella*-positive donor cells between the paired genotypes were equivalent within the same mouse (Fig. S2C). Therefore, non-cell autonomous mechanisms contribute to protection conferred by *Atg16L1^T316A^* heterozygosity.

*Atg16L1^A/A^* macrophages from human donors and knock-in mice display aberrant toll-like receptor (TLR)-NFkB signaling and overproduction of IL-1β, TNFα, and other cytokines (*8, 21, 33, 34*). To determine whether *Atg16L1^T316A^* heterozygosity alters the immune status of macrophages, we performed RNA-seq analysis of BMDMs from wild-type (*Atg16L1^T/T^*), *Atg16L1^T/A^*, or *Atg16L1^A/A^*mice two hours post-inoculation with *Salmonella* or mock treatment. Principal component analysis of the most variably expressed genes separated samples by treatment condition on PC1 and genotype by PC2 (Fig. 2A). Non-hierarchical clustering showed a pattern of gene up- and down-regulation in response to infection, with uninfected *Atg16L1^T/A^*or *Atg16L1^A/A^* clustering separately from wildtype but forming more distinct clusters in the infected group (Fig. S3A). In response to the infection, over 2000 of the significantly upregulated genes were shared among all three genotypes. However, over 900 genes upregulated were uniquely shared between *Atg16L1^T/A^* and *Atg16L1^A/A^* and not *Atg16L1^T/T^* (Fig. 2B), compared with only 94 and 67 that were shared between *Atg16L1^T/A^* and *Atg16L1^T/T^* or *Atg16L1^A/A^*and *Atg16L1^T/T^*, respectively. These results demonstrate that one copy of *Atg16L1^T316A^* recapitulates a large proportion of transcriptomic shifts that occur in homozygote cells.

**Fig. 2.**
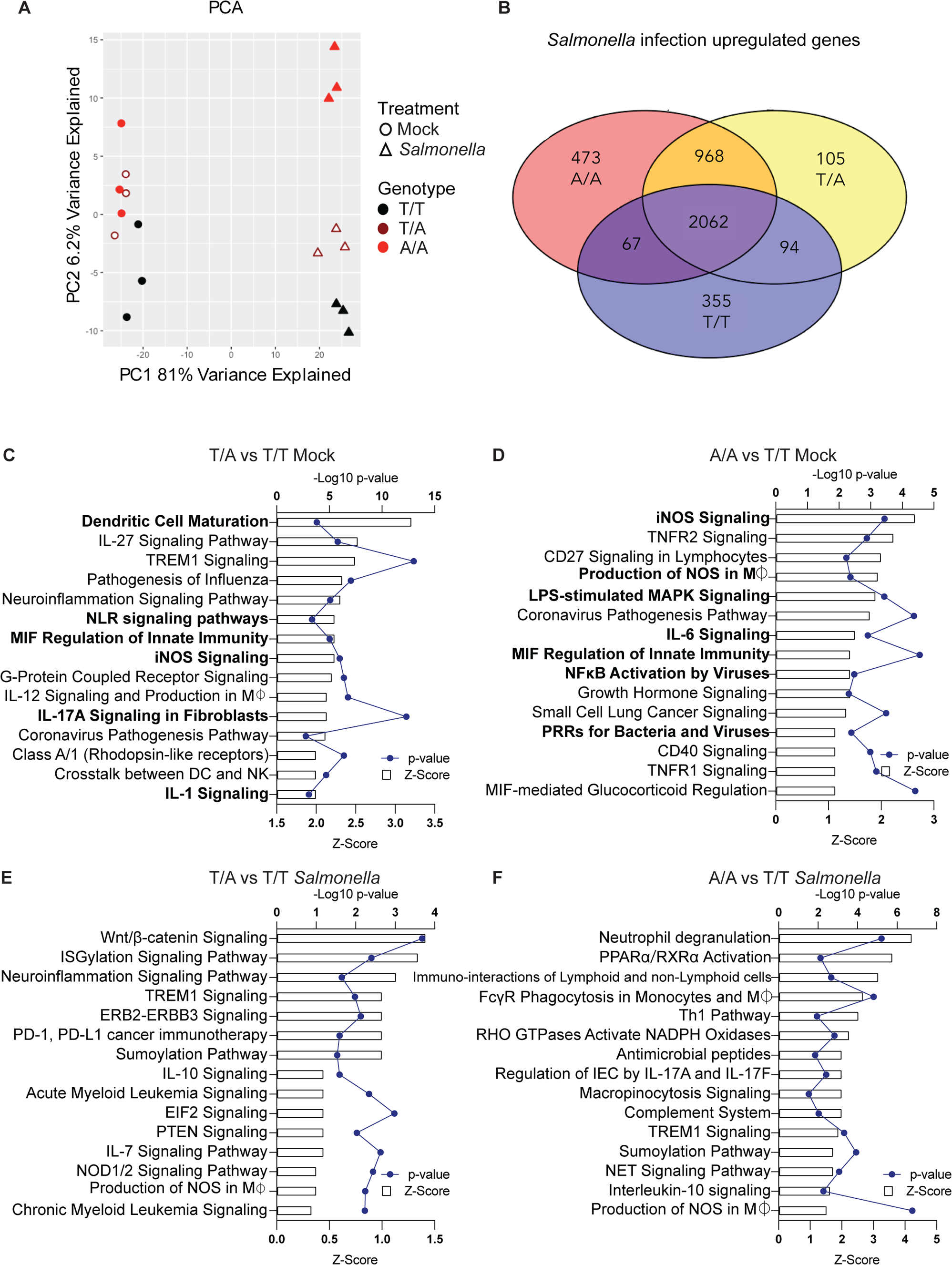
One copy of *Atg16L1 T316A* is sufficient to mediate a proinflammatory gene expression signature. (**A**) BMDMs (n=3 each condition) of indicated genotypes either mock treated or infected with *Salmonella* at multiplicity of infection (MOI) of 100 for 2 hours, then total RNA of these BMDMs were collected and subjected for bulk RNAseq. Principal Component Analysis (PCA) was done with 746 most variably expressed genes identified by DEseq across all the samples. (**B**) Venn diagram of genes upregulated by *Salmonella* infection in BMDMs of indicated genotypes. (**C-F**) Ingenuity Pathway Analysis of enriched pathways and the corresponding p-values + Z-Scores were listed for comparisons between indicated genotypes under either mock condition (C-D) or *Salmonella* infected condition (E-F). Select pathways associated with innate immunity in mock-treated samples are indicated in bold.

Ingenuity Pathway analysis (supplementary table 1) showed that, even under mock treatment conditions, *Atg16L1^T/A^* BMDMs displayed increased innate immune activation compared to wildtype, including upregulation of genes associated with activating dendritic cells and macrophages, iNOS signaling, and the NLR-IL1-IL17 inflammatory axis (Fig. 2C). Mock-treated *Atg16L1^A/A^* BMDMs displayed similar enrichment of transcripts associated with innate immunity (Fig. 2D). Pathways that were differentially regulated when comparing mock-treated *Atg16L1^T/A^* and *Atg16L1^A/A^*BMDMs were not as obviously associated with innate immunity and had low z-scores (Fig. S3B). *Atg16L1^T/A^* and *Atg16L1^A/A^* BMDMs treated with *Salmonella* still shared enrichment of immune-associated transcripts when compared to wildtype, such as TREM1 signaling, ISGylation, IL-10 signaling, and NOS production (Fig. 2E-F). Despite these similarities between *Atg16L1^T/A^* and *Atg16L1^A/A^* BMDMs, an antiviral interferon (IFN) signature distinguished homozygous from heterozygous cells (Fig. S3C), suggesting homozygous cells might experience stronger autophagy inhibition under *Salmonella* challenge, which would increase IFN activation (*34–37*).

If the *Atg16L1^T316A^* allele is preferentially expressed over the wildtype allele in heterozygous cell, this might explain the dominance conferred by the risk allele that has been proposed previously (*21, 33*). However, the wildtype *Atg16L1* allele was expressed at higher levels than risk allele by 14% in heterozygous cells (Fig. S3D), irrespective of treatment. Taken together, these data suggest that one copy of *Atg16L1^T316A^* is sufficient to achieve a pro-inflammatory transcriptional state that overlaps with homozygous cells.

### *Atg16L1 T316A* heterozygosity is sufficient to enhance inflammatory cytokines production

To determine whether T316A heterozygosity affects immune responses *in vivo*, we analyzed a panel of cytokines in the serum from mice 7 days post inoculation, a time point when *Salmonella* burden was substantially reduced in *Atg16L1^T/A^* mice (Fig. 1B and Fig. S4A). With the exception of IL-1α, which was not detectable, and CCL2, which was similar between the three groups, *Atg16L1^T/A^* mice had elevated levels of all cytokines (Fig. 3A). *Atg16L1^A/A^* mice displayed a similar increase in cytokine levels, although IL-2 and IFNγ were not significantly different from *Atg16L1^T/T^* mice. To test whether enhanced cytokine production in *Atg16L1^T/A^* animals is biologically significant, we used a sterile sepsis model involving LPS challenge (*38*). *Atg16L1^T/A^*and *Atg16L1^A/A^* mice displayed accelerated lethality compared with *Atg16L1^T/T^* mice (Fig. 3B).

**Fig. 3.**
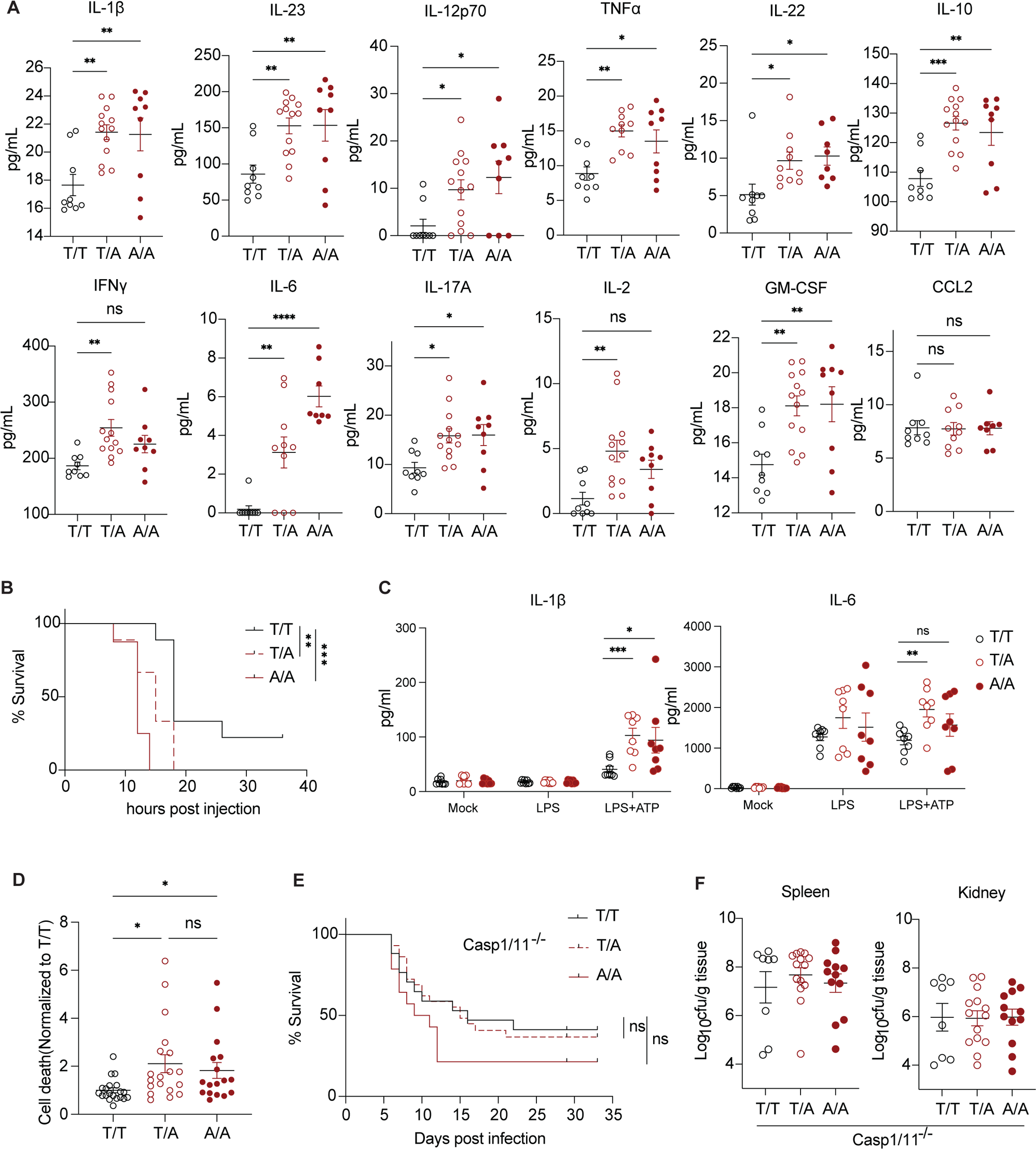
*Atg16L1 T316A* heterozygosity is sufficient to enhance cytokine production. (**A**) Multiplex bead array measurements of indicated cytokines in serum from mice infected with 2×10^4^ cfu *Salmonella* by oral gavage 7 days post infection. T/T, n=9; T/A, n=13; A/A, n=9. (**B**) Survival of mice injected with 27mg/kg LPS. T/T, n=9; T/A, n=9; A/A, n=8. (**C**) ELISA for IL-1β and IL-6 in supernatants from BMDMs of indicated genotypes that were mock treated or stimulated with 500ng/mL LPS for 5 hours alone or followed by 2mM ATP for 1 hour. (**D**) Relative cell death of peritoneal macrophages from mice injected IP with *Salmonella* from Fig. 1E measured by live/dead staining dyes and gating using flow cytometry. (**E**) Survival curve of *Atg16L1^T316A^* mice on the *caspase 1/11^−/−^*background orally inoculated with 2×10^5^ cfu *Salmonella*. T/T, n=17; T/A, n=28; A/A, n=14. (**F**) Bacterial burden in indicated organs of mice 2 days post IP inoculation with 2×10^5^ cfu *Salmonella*. Data points in (A, D and F) represent individual mice, in B are pooled from 3 independent experiments, and bars represent mean ± SEM. All data were collected from at least two independent experiments. ^∗^p < 0.05; ^∗∗^p < 0.01; ^∗∗∗^p < 0.001; ^∗∗∗∗^p < 0.0001; ns, not significant, by log-rank Mantel-Cox test in (C) and (E), one-way ANOVA with Holm-Sidak test for (A, B and D).

One of the first immune functions associated with ATG16L1 and autophagy is the suppression of inflammasomes, multiprotein innate immune complexes that activate caspase-1 to generate the mature form of IL-1β and associated cell death pyroptosis (*39, 40*). Inflammasome regulation by autophagy acts at multiple levels, including targeting inflammasome subunits such as ASC, NLRP3 and NLRP1 for degradation (*41, 42*), preventing accumulation of mitochondrial reactive oxygen species (ROS) (*43, 44*), and negative regulation of signaling components upstream of cytokine transcription (*34*). Both *Atg16L1^T/A^* and *Atg16L1^A/A^*BMDMs produced more IL-1β than wildtype cells when stimulated with LPS and ATP to trigger the NLRP3 inflammasome (Fig. 3C). T316A heterozygous cells also produced more IL-6 than wildtype. Considering that *Salmonella* can activate other inflammasomes in addition to NLRP3 (*45*), we subjected BMDMs to conditions that activate the NLRC4 and AIM2 inflammasomes but did not detect a significant effect of the T316A allele (Fig. S4B). Together, these results indicate that one copy of *Atg16L1^T316A^* is sufficient to enhance cytokine production and NLRP3 inflammasome activation.

Consistent with enhanced inflammasome activation *in vivo*, *Salmonella* infection in both *Atg16L1^T316A^* heterozygous and homozygous mice led to increased cell death of peritoneal macrophages compared with wildtype mice (Fig. 3D). To test whether the inflammasome is necessary for the enhanced protection, we crossed *Atg16L1^T316A^* mice to Caspase1/11 double knockout (*Casp1/11^−/−^*) animals, which are less susceptible to *Salmonella* than single Caspase-1 knockout mice allowing us to assess the effect of *Atg16L1* mutation(*46*)*. Atg16L1^T/A^*;*Casp1/11^−/−^* mice lost protection from lethality and displayed similar bacterial burden as *Atg16L1^T/T^*;*Casp1/11*^−/−^ and *Atg16L1^A/A^*;*Casp1/11^−/−^* mice (Fig. 3E-F). These results demonstrate that increased inflammasome activation contributes to protection conferred by *Atg16L1^T316A^* heterozygosity.

### *Atg16L1^T316A^* heterozygosity and homozygosity differentially affect LAP to impact *Salmonella* susceptibility

Although the increased inflammatory response can explain the protection conferred by *Atg16L1^T316A^* heterozygosity, it was unclear why *Atg16L1^T316A^* homozygosity was not protective given our results showing that having two copies of this variant also enhances cytokine production and immune signaling in mice and BMDMs. One possibility is that there is a second function of ATG16L1 (in addition to inhibiting cytokines) that is differentially affected by *Atg16L1^T316A^* heterozygosity and homozygosity. The amino acid substitution occurs adjacent to a WD40 repeat domain in the C-terminus that is necessary for CASM (*47*), and thus, autophagosome-independent functions may be sensitive to mutation in this position. CASM pathways are important in a variety of infection models (*48–51*). The best described version of CASM is LC3-associated phagocytosis (LAP), where phagosomes bearing internalized pathogens or their byproducts are decorated with LC3 to promote their fusion with the lysosome (*5, 52*). Rubcn, which is essential for LAP but dispensable for autophagy (*52*), promotes defense against *Salmonella* infection in zebrafish (*53*). Therefore, we analyzed the impact of *Atg16L1^T316A^* on LAP by performing an established assay in which phagocytic cells are incubated with fluorescent beads coated with the TLR2-agonist zymosan (*54*). As previously shown, BMDMs from wildtype *Atg16L1^T/T^* mice displayed an intense LC3+ halo surrounding zymosan beads at every time point examined with a peak at 40 minutes, while *Rubcn^−/−^* BMDMs showed background levels of association (Fig. 4A-B). Although *Atg16L1^T/A^* and *Atg16L1^T/T^* BMDMs were similar, *Atg16L1^A/A^* BMDMs resembled *Rubcn^−/−^* cells and lost the colocalization between LC3 and zymosan beads (Fig. 4A-B). We conclude that *Atg16L1^T316A^* homozygosity disrupts the LAP pathway in murine macrophages, but a single copy of the wildtype allele is sufficient to maintain LAP functionality.

**Fig. 4.**
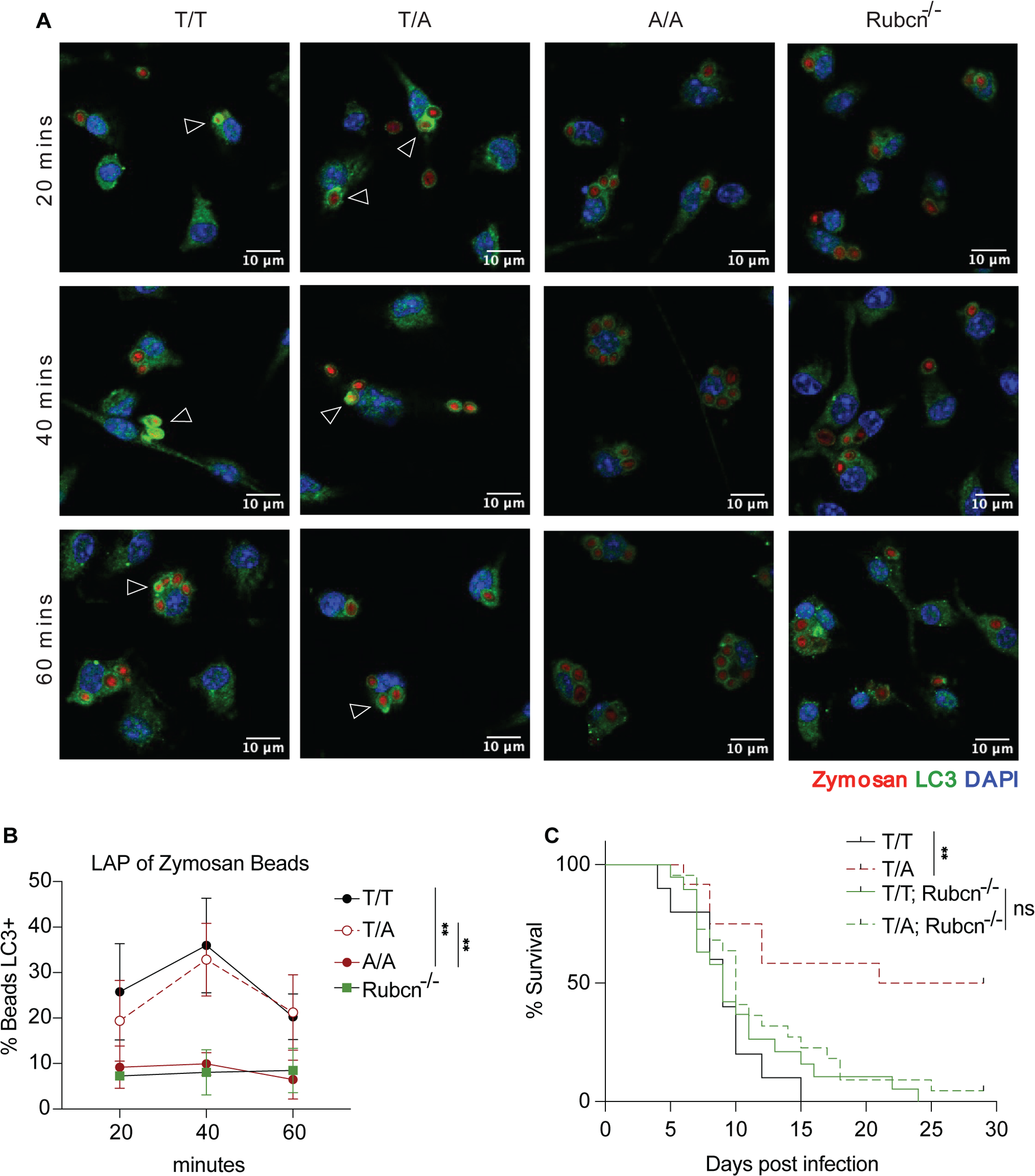
LAP is impaired by *Atg16L1 T316A* homozygosity and not heterozygosity. (**A**) LC3 staining of BMDMs from *Atg16L1^T316A^* knock-in or *Rubcn^−/−^* mice incubated with zymosan-conjugated fluorescent beads for the indicated time. DAPI shown in blue, Zymosan beads in red and LC3 staining in green. Arrows indicate LC3+ beads, images are representative of 3 independent experiments. (**B**) Quantitation of proportion of LC3+ beads from (A). (**C**) Survival curve of *Atg16L1^T316A^* knock-in mice on a *Rubcn^−/−^* background following oral inoculation with 2×10^4^ cfu *Salmonella*. *Atg16L1^T/T^* n=10 ; *Atg16L1^T/A^* n=12; *Atg16L1^T/T^* ;*Rubcn^−/−^* n=19; *Atg16L1^T/A^*;*Rubcn^−/−^* n=22. Data in B are mean ± SEM. All data were collected from at least two independent experiments. ^∗∗^p < 0.01; ns, not significant, by two-way ANOVA with Tukey’s multiple comparisons test for (B), and Log-rank Mantel-Cox test in (C),

To determine the role of LAP in protection mediated by *Atg16L1^T316A^* heterozygosity, we crossed *Rubcn^−/−^*mice with *Atg16L1^T/T^* or *Atg16L1^T/A^* animals. Although LAP-sufficient *Atg16L1^T316A^* heterozygous mice demonstrated the same protection we observed previously compared with wildtype mice, *Atg16L1^T/A^*;*Rubcn^−/−^* mice lost this protection and succumbed to infection similar to *Atg16L1^T/T^*;*Rubcn^−/−^* controls when challenged with *Salmonella* (Fig. 4C). These results indicate that LAP is required for heterozygote-mediated protection during *Salmonella* infection, and that *Atg16L1^T316A^* heterozygosity and homozygosity differentially affect LAP.

### *ATG16L1 T300A* heterozygosity is sufficient to enhance cytokine production and maintain LAP in human macrophages

We next asked if these observations in murine cells can be extended to humans. Several studies have reported increased inflammasome activity in cells from *ATG16L1^A/A^* donors, but samples from heterozygous individuals have not been assessed (*8, 55*). We generated macrophages derived from CD14+ monocytes (MDMs) from peripheral blood samples from 45 healthy individuals that were then stimulated with LPS and ATP. We assessed samples for the *ATG16L1^T300A^*allele after collecting cytokine data to ensure that these experiments were performed blind to the genotype. Our cohort was comprised of roughly equal numbers of each of the three *ATG16L1* genotypes (Fig. 5A), consistent with reported allele frequencies (*34*). *ATG16L1^T/A^*MDMs produced highest levels of IL-1β when compared to both *ATG16L1^T/T^*and *ATG16L1^A/A^* MDMs (Fig. 5B). We also measured secretion of IL-8 to assess the functionality of the TLR-MAPK-NFkB pathway (*56*). We did not observe enhanced production of this cytokine in either homozygous or heterozygous carriers of the risk allele (Fig. 5C). MDMs from *ATG16L1^T/T^*and *ATG16L1^T/A^* donors displayed similar LC3 recruitment to internalized zymosan beads, while those from *ATG16L1^A/A^* carriers did not display colocalization above background levels (Fig. 5D-E). We conclude that in humans, similar to what we discovered in mice, one copy of the T300A allele is sufficient to drive inflammasome hyperactivity, while two copies are required to impair LAP.

**Fig. 5.**
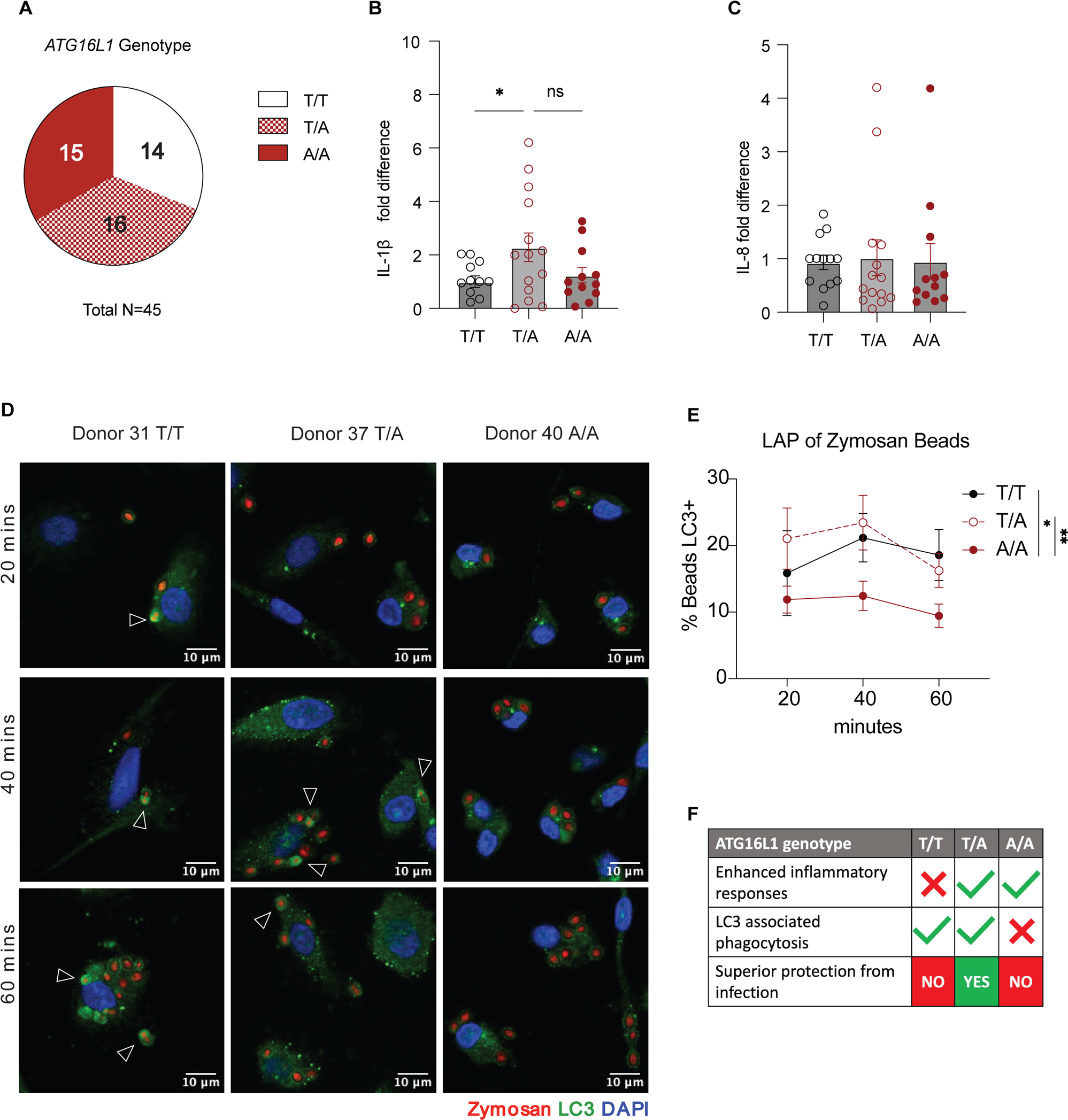
*ATG16L1 T300A* heterozygosity is sufficient to enhance cytokine production and maintain LAP in human macrophages. (**A**) Frequency of T300A genotypes among donor group. *ATG16L1^T/T^* n=14; *ATG16L1^T/A^*n=16; *ATG16L1^A/A^* n=15. (**B**) IL-1β secretion by monocyte-derived macrophages (MDMs) from human donors. Cells were stimulated with 500ng/mL LPS for 5 hours followed by 30 minutes 5mM ATP pulse. Summary of 5 individual experiments, measurements from each were normalized to non-risk average from that batch. (**C**) IL-8 secretion by MDMs from human donors treated as in B. Summary of 5 individual experiments, measurements from each were normalized to non-risk average from that batch. (**D**) MDMs from human donors were treated with Zymosan-conjugated fluorescent beads for the indicated time and stained for endogenous LC3. DAPI shown in blue, Zymosan beads in red and LC3 staining in green. Arrows indicate LC3+ beads, images are representative of 3 independent experiments.(**E**) Quantitation of images in D. *ATG16L1^T/T^*n=4; *ATG16L1^T/A^* n=5; *ATG16L1^A/A^* n=7. (**F**) Model: superior defense against bacterial infections requires both enhanced inflammatory responses, conferred by the 300A allele of ATG16L1, and intact LC3 associated phagocytosis, which requires at least one copy of the 300T allele. Data in (B, C and E) are mean ± SEM. All data were collected from at least two independent experiments. ^∗^p < 0.05; ^∗∗^p < 0.01; ns, not significant, by one-way ANOVA with Fisher’s LSD test for (B) and two-way ANOVA with Tukey’s multiple comparisons test for (E).

## Discussion

We identified a beneficial effect of the common *ATG16L1^T300A^* Crohn’s disease risk allele and its murine equivalent during bacterial infections, which was exclusive to heterozygosity. One copy of *ATG16L1^T300A^* was sufficient to enhance cytokine production and inflammasome activation, which was necessary for protection. Although *Atg16L1^T300A^*homozygosity also enhanced cytokine production, having two copies inhibited LAP, a process that remained intact in *Atg16L1^T300A^* heterozygotes. Thus, *Atg16L1^T300A^* heterozygosity selectively impacts one function of the protein, cytokine inhibition, while *Atg16L1^T300A^* homozygosity inhibits a second critical antimicrobial function, LAP. This decoupling of ATG16L1 functions explains why *Atg16L1^T316A^* homozygous mice were unable to take advantage of the enhanced cytokine production that protects against *Salmonella* (Fig. 5F).

Our results are in line with prior studies showing that *ATG16L1^T300A^* is a conditional loss of function variant. In the intestinal epithelium, the cell-autonomous function of autophagy in Paneth cells is severely impacted by *ATG16L1^T300A^*, which reflects the high secretory burden and dependence on organelle homeostasis for viability of these cells (*9, 36, 57–59*). The role of ATG16L1 in the epithelium may be particularly important early during the infection prior to dissemination(*17, 57*). Also, the reason the inflammasome- and cytokine-suppressing function of ATG16L1 is prominent in the setting of our experiments may be because *Salmonella* (and *L. monocytogenes*) evade cell-autonomous inhibition through autophagy, which would be consistent with our finding that *Atg16L1^T/A^* and *Atg16L1^A/A^* macrophages display similar intracellular bacterial burden *in vitro* and in the mixed BM chimeras *in vivo*. Although several mechanisms by which T300A impairs ATG16L1 function have been proposed (*8, 9, 21*), we suggest these are not mutually exclusive if autophagy and CASM functions of ATG16L1 are differentially affected by T300A, and the exact impact of the substitution depends on the cell type or environmental stressor. Our results are reminiscent of protection against malaria and trypanosomiasis conferred by heterozygosity for the sickle cell anemia allele of *HBB* and kidney disease allele of *APOL1*, respectively. Balancing selection may occur in these examples because genetic diseases preferentially develop in homozygotes, but the protection against a life-threatening parasite infection conferred by one copy of these disease risk alleles provides a substantial advantage (*60, 61*). We speculate that the enhanced immunity provided by the CD risk allele of *ATG16L1* may explain its widespread prevalence.

## Acknowledgments

We would like to acknowledge New York University Grossman School of Medicine Reagent Preparation, Flow Cytometry and Cell Sorting, Microscopy, Genome Technology laboratories for use of their instruments and technical assistance. Core facilities were supported by NIH grant P31CA016087.

## Funding

National Institutes of Health grant R01 AI121244 (KC, VJT)

National Institutes of Health grant R01 DK093668 (KC)

National Institutes of Health grant R01 HL123340 (KC)

National Institutes of Health grant R01 AI140754 (BS, KC, VJT)

Crohn’s & Colitis Foundation (KC)

F31 HL149238 (ER)

F31 DK111139 (PKM)

## Author contributions

XY and ER were the key contributors in designing and conducting most of the experiments. PKM generated the initial observations that established the study. XY, ER, PKM, VJT, BS, KC conceived of and directed the project. XY, ER, and KC wrote the manuscript. EZ and SAL assisted with key experiments. All authors commented on and edited the manuscript.

## Data and materials availability

RNA-seq data discussed in the study have been deposited in the Gene Expression Omnibus with an access number of GSE255216. The biological material is available upon request to KC. All data are available in the main text or the supplementary materials.

## Materials and Methods

### Mice

All mice used were age and sex matched 8-14-week-old C57BL6 mice with littermate controls. *Atg16L1^T316A^* mice were previously described (*8*). *CAG-EGFP* (003291), *Cd45.1* (002014), and *Caspase1/11^−/−^* (016621) mice were purchased from Jackson Laboratory and *Rubcn*^−/−^ were generously provided by Dr. Doug Green and previously described, and these mice were bred onsite. Mice crossed to *Atg16L1^T316A^* mice to generate double mutants were bred to fix the knockout allele first and maintained as T316A heterozygotes to generate littermates of all three genotypes (wildtype, heterozygous, and homozygous for *Atg16L1^T316A^*) for experiments. The *Atg16L1^T316A^* mouse line used in Figure 1I was generated by using the CRISPR ribonucleoprotein (RNP) system containing a chemically-modified crRNA + tracrRNA complexed with Cas9 protein (ctRNP). All reagents were obtained from from IDT. A single-stranded synthetic oligodeoxynucleotide (ssODN) repair template containing the T300A coding DNA sequence *gct* replacing the wild-type *act* sequence at Exon 9, was microinjected along with the ctRNP into C57BL/6 zygotes to generate mice (*62*). The T316A mutation was verified by Sanger sequencing in the resulting gene-modified mice. All animal studies were performed according to approved protocols by the NYU School of Medicine Institutional Animal Care and Use Committee (IACUC).

### *S.* Typhimurium and *L. monocytogenes* infections

For both bacteria, single colonies were selected. *S.* Typhimurium strain SL1344 was grown overnight in Luria-Bertani broth with 10ug/mL streptomycin with shaking at 37°C. *L. monocytogenes* strain 10403s was grown overnight in brain heart infusion (BHI) broth with 5ug/mL streptomycin with shaking at 37°C. Bacterial suspension was diluted 1:100 followed by an additional 2-3 hours of growth until bacteria were at an optical density of 0.8-1.2 at 600nm. Bacterial density was confirmed by dilution plating. For oral *S.* Typhimurium infection, mice were inoculated by gavage with 2 × 10^4^ cfu or 2 × 10^5^ cfu resuspended in 100 μl PBS. Mice were infected intraperitoneally (IP) with 2 × 10^5^ cfu resuspended in 100 μl PBS. For *L. monocytogenes* infection, mice were inoculated IP with 3 × 10^5^ cfu resuspended in 100 μl PBS. Survival was monitored over time. For quantification of bacterial shedding in stool and bacterial dissemination to liver, kidney, and spleen, samples from individual mice were weighed, homogenized in PBS 0.5% Triton X-100, and plated in serial dilutions on LB or BHI agar streptomycin plates. *In vitro* infection was carried out by opsonizing bacteria grown as described above with normal mouse serum followed by washing and resuspension in PBS. Cells were inoculated at desired MOI followed by centrifugation at 300g x 5’ incubation for 30 minutes at 37C. Excess bacteria was then washed off and cells were incubated for the remaining time with 50µg/mL gentamycin (Sigma).

### Bone marrow chimera

6-week-old recipient mice on C57BL6 background, either CD45.1 congenic mice or GFP+ mice, were lethally irradiated with two doses of 500 rads with a 4-hour interval, followed by retroorbital injection with 2×10^6^ donor bone marrow cells to allow reconstitution. 8 weeks later, mice were infected with *Salmonella* as indicated. Upon sacrifice, the bone marrow reconstitution rates were determined by flowcytometry by looking at the relative abundance of donor vs recipient hematopoietic cells.

### Flow cytometry

To quantify intracellular *S.* Typhimurium infection, mice were infected with mKO-expressing *Salmonella* generously provided by Dr. Brett Finlay (*63*) which is detectable in the PE channel. Peritoneal exudate cells were isolated by lavage with 8mL ice-cold PBS. Cells were then washed and treated with 150,000U lysozyme at RT for 15 minutes to remove extracellular bacteria. Cells were then stained for surface markers: anti-mouse CD45 (clone 30-F11) or anti-mouse CD45.1 (clone A20) and anti-mouse CD45.2 (clone 104), Dump staining (anti-mouse CD3e (clone 145-2C11), anti-mouse CD19 (clone 6D5) and anti-mouse NKp46 (clone 29A1.4)), anti-mouse CD11b (clone M1/70), anti-mouse Ly6G (clone 1A8) or anti-mouse Gr1 (clone RB6-8C5), anti-mouse Ly6C (clone HK1.4), and anti-mouse F4/80 (clone BM8). Fixable LIVE/DEAD stain (Invitrogen) was used to exclude dead cells. And CD45+ (or either CD45.2+ or CD45.1+) Dump-cells are further gated to define macrophages as Cd11b+, Ly6G-Ly6C- (or Gr1-) and F4/80+; while neutrophils were defined as CD11b+, Gr1+ (or Ly6G+Ly6C-) and F4/80-. Salmonella containing cells were defined as mKO+ cells. Dead peritoneal cells were quantified by either DAPI or Live/Dead staining.

### BM-derived Macrophage Isolation and Culture

BM cells from mouse femurs and tibia were collected by centrifugation, followed by red blood cell lysis in ACK buffer and resuspension in DMEM. Cells were cultured for 3 days on non-treated plates in DMEM supplemented with L-Glutamine, NEAA, Sodium Pyruvate, B-mercaptoethanol and Pen/Strep in the presence of 100ng/ml M-CSF (576406, Biolegend) followed by 3 more days with renewed media and cytokines. Differentiated macrophages (BMDMs) were then collected and plated for stimulation or microscopy.

### LC3-Associated Phagocytosis (LAP) Assay

BMDMs and human macrophages were plated on glass coverslips in a 24-well plate and treated with 1.25ug/mL Zymosan-coated Texas Red ^TM^ fluorescent beads (Thermo Z2843) at 37C for indicated times. Cells were then washed with cold PBS, fixed with 4% PFA followed by permeabilization in 0.2% Triton X-100 PBS and blocking with 0.1% Triton X-100 2.5% BSA 5% Newborn goat serum. Cells were then stained with primary antibody rabbit anti-mouse LC3 (PD014, MBL, 1:200 in blocking buffer) and fluorescent (AF488) conjugated goat anti-rabbit secondary antibody (Thermo, 1:1000 in blocking buffer). Cells were then washed in PBST and sealed with DAPI containing mounting medium (Dapi-Fluoromount-G™, Electron Microscopy Sciences). Images were taken by Zeiss 880 confocal microscopy. LC3-positive beads were identified using FIJI software by measuring the ratio of the mean LC3 fluorescence associated with Zymosan beads over cytosolic background signal in the same cell, and threshold for LAP activity was determined by comparing wildtype cells to *Rubcn^−/−^*.

### Human peripheral blood-derived macrophages

LeukoPaks were obtained from anonymous blood donors with informed consent from the New York Blood Center. Peripheral blood mononuclear cells (PBMCs) were isolated and differentiated into macrophages for experimentation as previously described (*64, 65*). Briefly, CD14+ monocytes were isolated from PBMCs using the EasySep Human CD14 Positive Selection Kit (Stem Cell) and cultured with 100U/mL GM-CSF (Peprotech) for 4 days prior to plating for stimulation. After replating, PMA (100ng/ml) was added to differentiate macrophages for 48hr, then incubated with fresh culture medium to rest overnight prior to experimentation.

### RNA isolation and qPCR

Cells were collected and resuspended in 700uL of RLT buffer (Mini RNeasy Kit) and homogenized using TissueRuptor (Qiagen). RNA was then isolated using the RNeasy Mini Kit (Qiagen) as per manufacturer’s instructions, DNase treatment was performed using RNeasy DNase kit (Qiagen) and protocol. cDNA synthesis was performed using ProtoScript M-MuLV First Strand cDNA synthesis kit (New England Biolabs) and random hexamer primers. qPCR was performed on a Roche480II Lightcycler.

### Cytokine quantification

Individual cytokines in supernatants from cell culture experiments were measured using the species-appropriate ELISA Kits mouse IL-6 (Invitrogen™ 88706488), mouse IL-1beta (Invitrogen™ 88701388), human IL-8 (Invitrogen™ 88808677), human IL-1beta (Invitrogen™ 88726177) according to the manufacturer’s instructions. For multiplexed measurement of serum cytokines, blood was collected via cheek bleed form *S.* Typhimurium infected mice. Sera was isolated spinning blood at 3000rpm, 10min, RT. The LEGENDplex mouse inflammation panel was used to quantify cytokines by flow cytometry (Biolgened cat#740150) according to the manufactural instructions.

### RNA sequencing

BMDMs were infected with late log phase *S.* Typhimurium at MOI 100 for 2hr. BMDM RNA was collected as indicated above for qPCR and cDNA libraries were prepared with the Ovation Ultralow RNA-seq System V2 (NuGEN) and sequenced on Illumina’s HIseq2 platform using a paired end protocol at the NYU Genomics Core. Reads that passed FASTQC were aligned to the most recent UCSC mouse genome assembly using STAR. Raw transcript counts were extracted by FeatureCount, DEseq2 was used to analyze differential gene expression. PCA plot was generated using DESeq2 with the 95 percentile most variable genes as input. Heat map was generated in DESeq2 including genes with absolute Log2 fold change > 1 and adjusted p value < 0 .01. Gene ontology pathway analysis was performed with IPA (QIAGEN), by including genes with adjusted p-value < 0.05 and absolute Log2 fold change > 1.

### Statistical analysis

All analyses except for RNA–seq data used Graphpad Prism v.10. An unpaired two-tailed *t*-test was used to evaluate differences between two groups where data was distributed normally with equal variance between conditions. An ANOVA with Holm–Sidak multiple comparisons test was used to evaluate experiments involving multiple groups. The log-rank Mantel–Cox test was used for comparison of mortality curves. Other tests used were specified in figure legends.

### Data availability

The data that support the findings of this study are available from the corresponding author upon request. FASTQ files and normalized count table corresponding to the RNA–seq data have been deposited in a public database (Accession number GSE255216).

---

**Figure S1.**
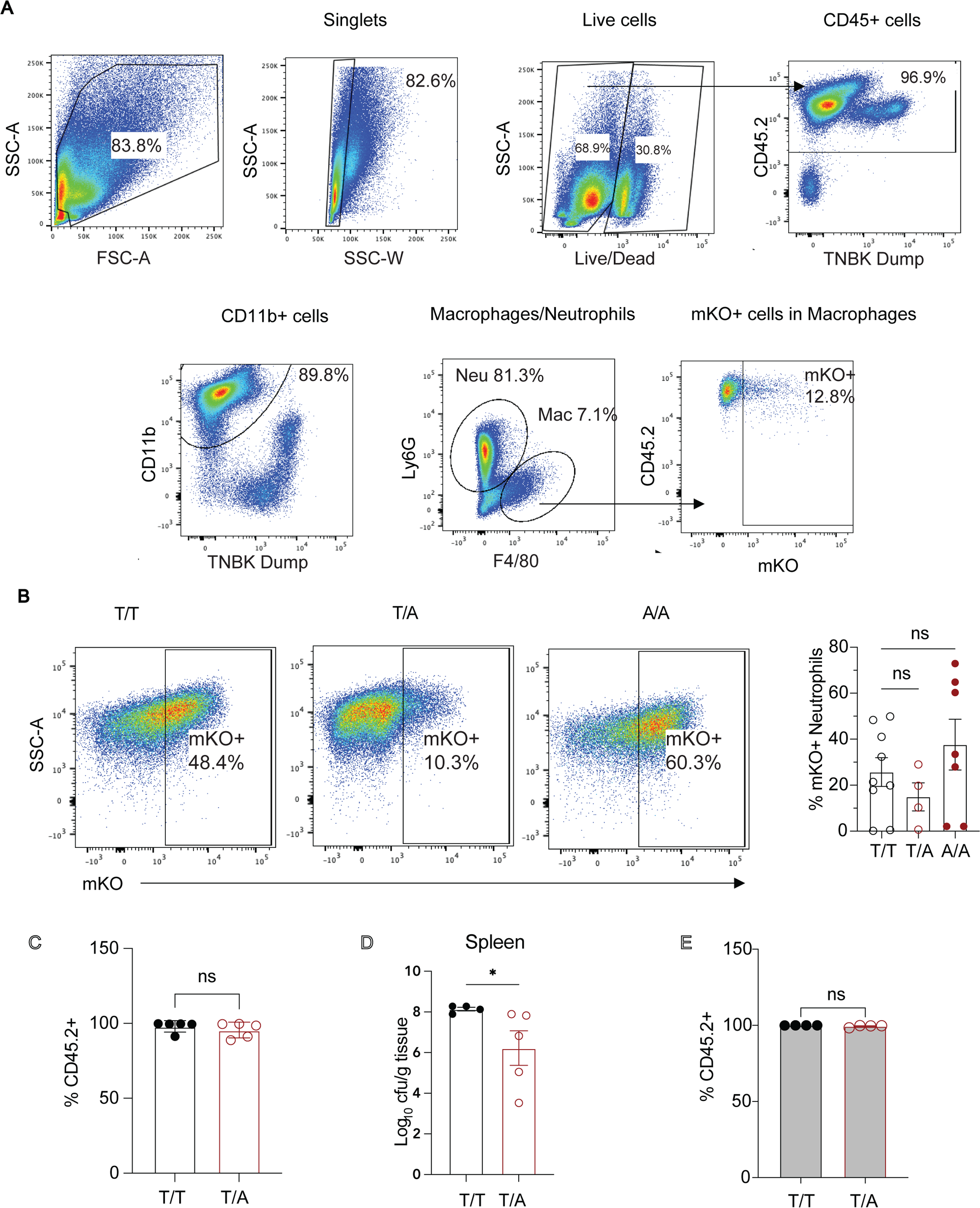
Flow cytometry data associated with Figure 1. A. Representative gating strategy for flow cytometry analysis of mKO+ peritoneal exudate cells 2 days following IP injection with mKO-expressing *Salmonella*. B. Proportion of mKO+ neutrophils among peritoneal exudate cells from mice following IP injection with mKO-expressing *Salmonella* from Figure 1D. Gating on live, CD45^+^, Ly6G^+^, CD11b^+^ cells for neutrophils. Atg16L1^T/T^ n=9; Atg16L1^T/A^ n=4; Atg16L1^A/A^ n=7. C. Chimeras generated by reconstituting CD45.1 mice with bone marrow of indicated genotypes, bone marrow reconstitution rate was examined by measuring CD45.2+ cells percentage in the peritoneal hematopoietic cells. D. Chimeras in C were infected as in Figure 1D and bacterial burden in spleen was determined 2 days post infection, n=5 each. E. Bone marrow reconstitution rate was examined in the bone marrow chimera experiments in Figure 1H by measuring CD45.2+ cells percentage in the peritoneal hematopoietic cells.

**Figure S2.**
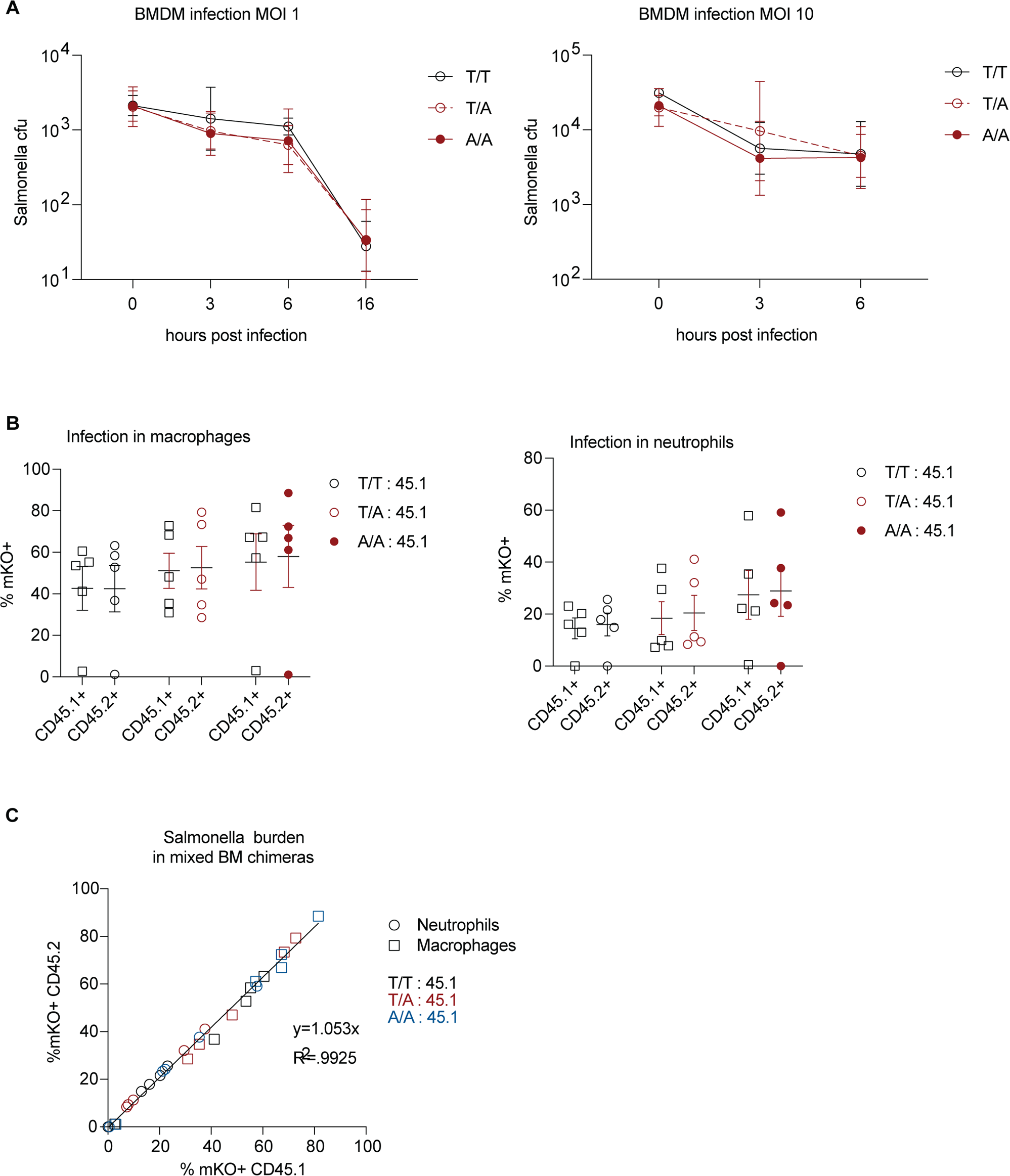
*Atg16L1^T316A^* heterozygosity does not enhance cell autonomous control of *Salmonella*. A. Quantification of intracellular bacteria in BM-derived macrophages (BMDMs) from *Atg16L1^T/T^*, *Atg16L1^T/A^*, and *Atg16L1^A/A^* mice incubated with Salmonella at multiplicity of infection (MOI) of 1 or 10 over time. B. Lethally-irradiated CD45.2+ GFP+ mice were injected with CD45.2+ BM cells from *Atg16L1^T/T^*, *Atg16L1^T/A^*, or *Atg16L1^A/A^* mice mixed 1:1 with CD45.1+ congenic BM cells. After 6 weeks, mice were injected IP with mKO expressing Salmonella and the proportion of mKO+ peritoneal macrophages and neutrophils were determined by flowcytometry after gating on CD45.2+GFP- (donor *Atg16L1^T/T^*, *Atg16L1^T/A^*, or *Atg16L1^A/A^*) and CD45.1+GFP- (donor wild-type) cells. C. Correlation between proportion of mKO+ CD45.2+ and CD45.1+ peritoneal cells within each individual mouse.

**Figure S3.**
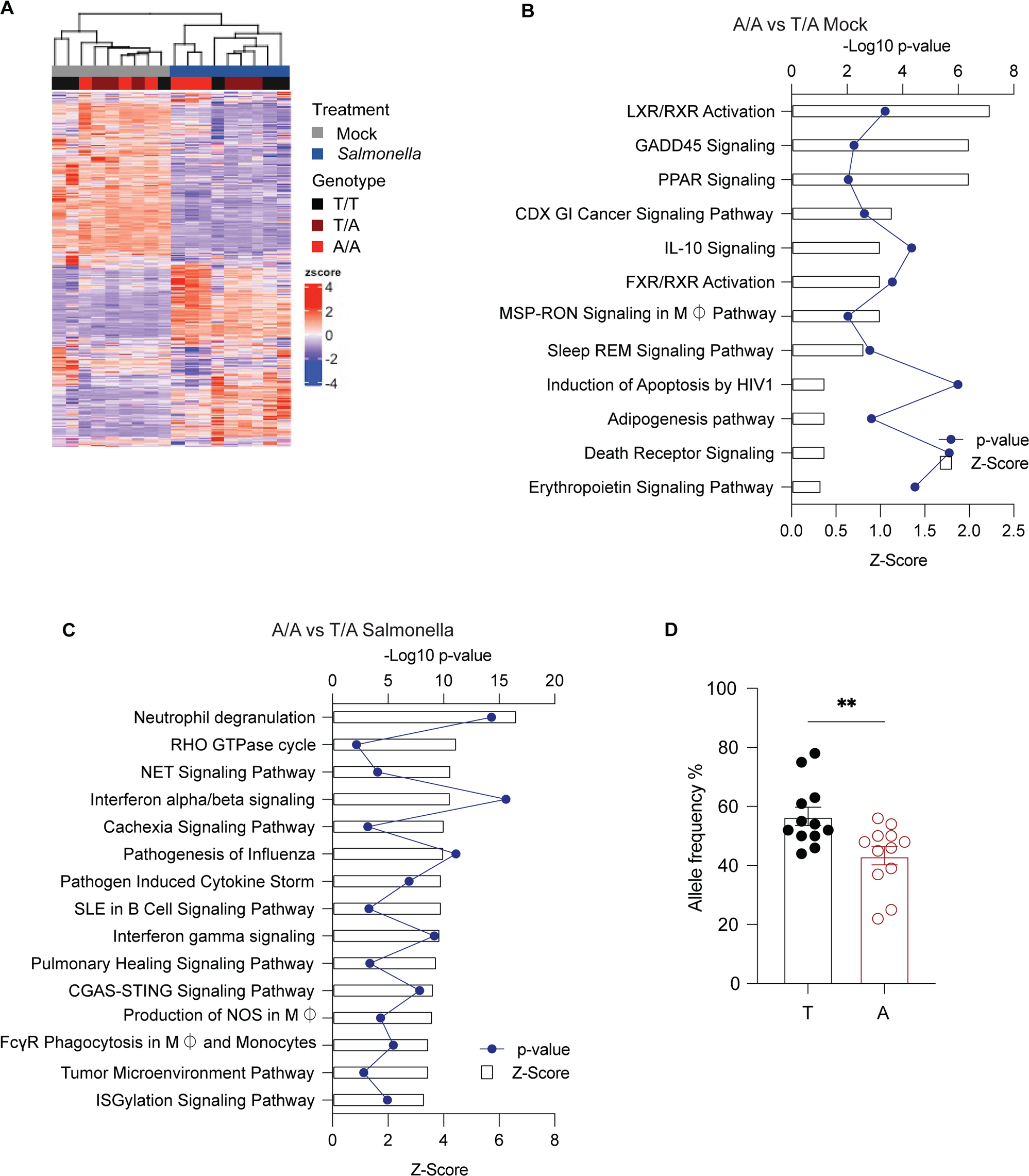
*Atg16L1^T316A^* impacts transcriptional responses in macrophages. A. Hierarchical clustering of differentially expressed genes in mock or Salmonella treated *Atg16L1^T/T^*, *Atg16L1^T/A^*, and *Atg16L1^A/A^* BMDMs. Color indicates row z-score, fold change > 2; FDR adjusted p-value < 0.01. B. Ingenuity Pathway Analysis of enriched pathways and the corresponding p-values + Z-Scores were listed for comparisons between A/A and T/A under mock condition. C. Ingenuity Pathway Analysis of enriched pathways and the corresponding p-values + Z-Scores were listed for comparisons between A/A and T/A under *Salmonella* infected condition. D. *Atg16l1^T300^* (T) vs *Atg16l1^A300^* (A) allele expression levels were examined in the heterozygous BMDMs. \*\**p*<0.01, two-tailed t-test.

**Figure S4.**
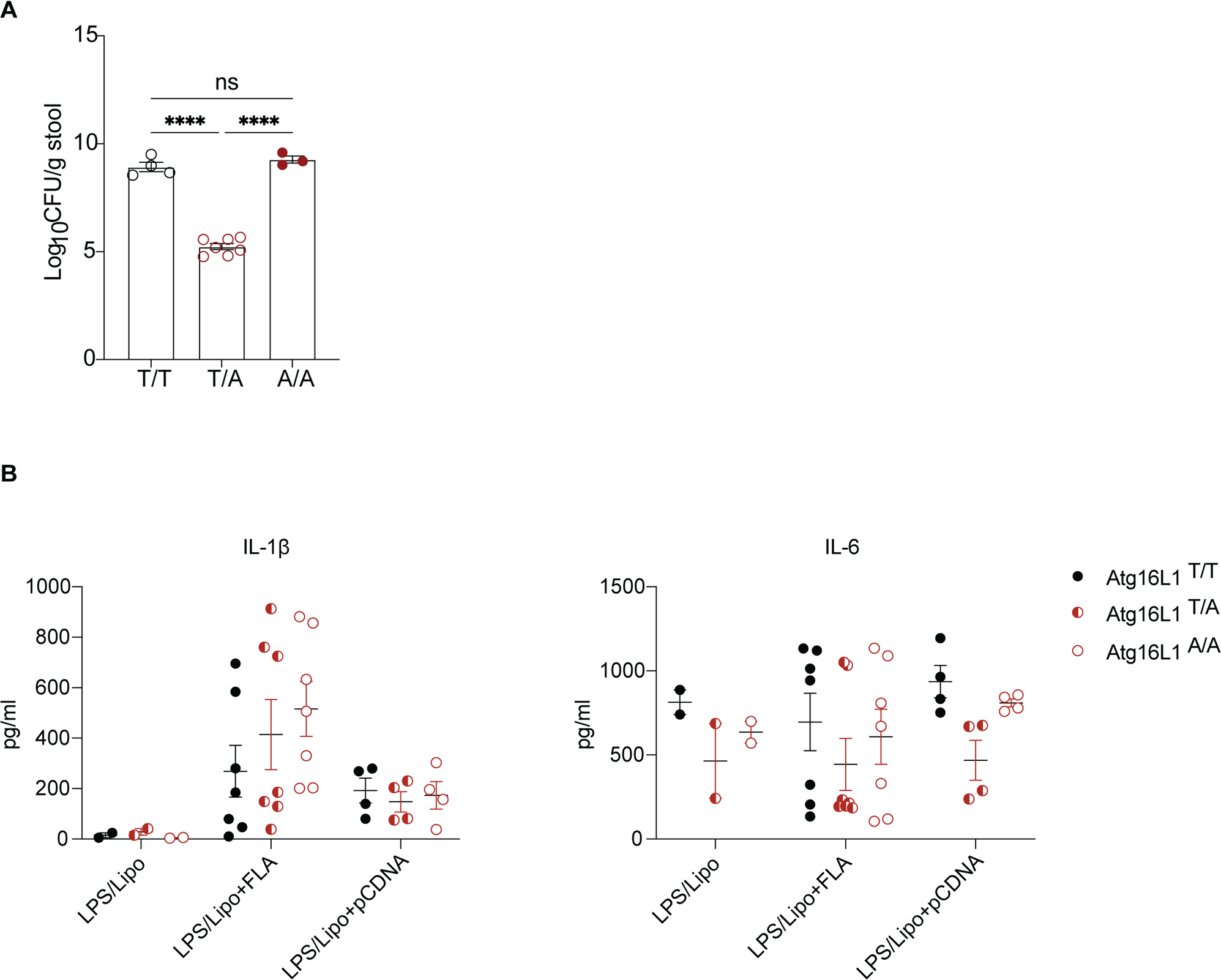
Effect of *Atg16L1 T316A* on cytokine production. A. Mice stool bacterial loads on day7 in Fig. 3A. B. Quantification of IL-1β and IL-6 by ELISA in supernatant from BMDMs primed with 500ng/ml LPS and stimulated with liposome (DOTAP) delivered flagellin (FLA) or vector plasmid DNA (pCDNA).

